# The extended ‘common cause’: causal links between punctuated evolution and sedimentary processes

**DOI:** 10.1101/2024.11.13.623443

**Authors:** P. David Polly

## Abstract

The common-cause hypothesis says that factors regulating the sedimentary record also exert macroevolutionary controls on speciation, extinction, and biodiversity. I show through computational modeling that common cause factors can, in principle, also control microevolutionary processes of trait evolution. Using Bermuda and its endemic land snail *Poecilozonites*, I show that the glacial-interglacial sea level cycles that toggle local sedimentation between slow pedogenesis and rapid eolian accumulation could also toggle evolution rates between long slow phases associated with large geographic ranges and short rapid phases associated with small, fragmented ranges and “genetic surfing” events. Patterns produced by this spatially driven process are similar to the punctuated equilibria patterns that Gould inferred from the fossil record of Bermuda, but without speciation or true stasis. Rather, the dynamics of this modeled system mimic a two-rate Brownian motion process (even though the rate parameter is technically constant) in which the contrast in rate and duration of the phases makes the slower one appear static. The link between sedimentation and microevolution in this model is based on a sediment-starved island system, but the principles may apply to any system where physical processes jointly control the areal extents of sedimentary regimes and species distributions.

**Non-Technical Summary:** The history of life is known from the fossils preserved in the geological record. The common-cause hypothesis suggests that processes like mountain building and sea-level change can affect both the structure of the geological record and species diversity. Using the snails of Bermuda as an example, this paper develops a computational model to show that sea level cycles could affect morphological evolution within species, not just species diversity. As sea level rose on Bermuda, the available snail habitat would have become smaller and more fragmented, which would be expected to drive rapid bursts of genetic drift (the random component of evolutionary change). When sea level fell, the snails’ habitat would have expanded and coalesced, resulting in slower rates of evolution because the total population size would increase. This process would produce an uneven cycle of rapid and slow evolution similar to what paleontologist Steven Gould observed in the fossil snails of Bermuda and led him to propose the theory of punctuated equilibria. While these simulations are focused specifically on an island system, the principles are applicable to other situations suggesting that geological and evolutionary processes may be linked in more ways than previously understood.

## Introduction

The non-random influences of sedimentary and stratigraphic processes on the fossil record are well known. Sedimentation rates, facies distributions, and erosion patterns are functions of sea level, accommodation space, and sediment influx (e.g., Catuneanu 2006). Ultimately, these non-random patterns in the stratigraphic record are driven by tectonic, eustatic, and climatic processes and the fossils preserved in the sediments inherit the imprint of the same stratigraphic controls (McKinney 1985; Kidwell 1986; Holland 2000; Smith et al. 2001; Bush et al. 2002; Kidwell and Holland 2002; Hannisdal 2006; Patzkowsky and Holland 2012). Paleontologists must account for this non-random to accurately interpret evolutionary and biodiversity patterns in the fossil record.

In this paper, I will show that Earth system processes can also influence microevolution evolution processes themselves. I will use computational modelling to show how sea-level cycles can simultaneously regulate rates of genetic drift (the stochastic component of evolutionary change) and the accumulation of stratigraphic sequences. My model is based on the rock and fossil records of Bermuda. That island’s sediment-starved and tectonically stable setting creates unusually clear links between sedimentary deposition and sea level cycles depending on whether the island platform is flooded or not. The model will show that genetic drift is very slow during low stands when the entire platform is subaerially exposed. Punctuated bursts of random change occur during high stands when small populations differentiate on isolated islets and the early falling-stage when ‘genetic surfing’ amplifies those differences by non-random sampling along the edges of their now-expanding geographic ranges (Excoffier and Ray 2008). The literal rises and falls of sea level force the rearrangement of the geographic ranges of island species in such a way that their rate of genetic drift is altered by the same processes that change the rate of sediment accumulation in which fossils are preserved. Another reason for focusing on Bermuda is that its endemic land snails (genus *Poecilozonites*) make up the clade that inspired Gould’s contribution to the theory of ‘punctuated equilibria’. While the punctuated equilibria model was based on principles of speciation by peripheral isolation (Eldredge and Gould 1972; Gould and Eldredge 1977), this paper will show that similar punctuated evolutionary patterns can, in principle, arise randomly from eustatically driven changes in a species geographic range without speciation or true stasis.

This paper focuses on the genetic drift component of evolution. Evolution is the net change in the mean values of a population’s (or species’) traits. Selection is the non-random component that arises from fitness differences in the traits, whereas genetic drift is the random component that arises from stochastic sampling of each generation from its progenitor (Wright 1931). Drift is a type of Brownian motion and is strongest when populations are small and sampling error is large. Drift is *always* a component of evolutionary change, whereas selection may be strong or weak, directional or stabilizing, or even absent depending on the context (Wright 1931; Lande 1976). Some authors have argued that true genetic drift is the dominant form of evolution (Kimura 1983) and others have found that Brownian motion patterns of trait evolution are fairly common in the fossil record (Hunt 2007), although one should note that Brownian motion can be produced by randomly fluctuating selection and that many authors have argued that stabilizing selection is the most common mode of phenotypic evolution (Polly 2004; Estes and Arnold 2007; Hunt 2007). Regardless of its historical role in shaping the evolutionary history of life, this paper focuses on genetic drift because its magnitude depends on net effective population size, which is related to the size and continuity of a species’ geographic range, which can be governed by the same factors that control sedimentary processes.

Why would sea level affect the rate of drift? The expected rate of genetic drift determined by the amount of genetic variance in the trait (*G*) relative to population size (*N*). If the genetic variance equals 1 unit (e.g., mm) and population size is 1000 then the rate of drift in the trait mean is 0.001 units per generation, but if population size drops to 10 then the rate of drift rises to 0.1. Drift is therefore faster in small populations, which makes it an important factor of evolution at times when populations sizes are small, in which case it can exceed the rate of evolution due to selection, or when selection is weak or absent (Lande 1976). Despite drift’s statistical simplicity, its real-world behavior can be complex and counter-intuitive because spatial processes can isolate or intermingle the local populations that make up a species and thus change the net population size, or can drive range expansions that result in non-random sampling of a single progenitor population so as to clone its traits across a large part of the species’ distribution (Ibrahim et al. 1996; Excoffier and Ray 2008; Polly 2019a). On islands, as this paper will show, sea-level cycles can drive cyclic patterns of fragmentation into small, isolated populations followed by range expansions into large panmictic ones. These processes can produce punctuated changes in rates of drift that mimic the patterns expected from the classic punctuated equilibria model of speciation. My goal is not to argue that Bermudian snails evolved solely by drift, but to demonstrate that complex, punctuated patterns of evolution can arise as a stochastic byproduct of the same factors that shape the stratigraphic record.

Why did sea-level change produce cyclic changes in sedimentation rate on Bermuda? Bermuda is a carbonate-topped seamount in the mid North Atlantic (**Fig. 1A**). It originated as a seafloor volcano along the North Atlantic ridge that was last active around the end of the Eocene (Reynolds and Aumento 1974). In its history, the seamount grew to as much as 3500 m above sea level before subsiding to become capped by a carbonate platform. The surface carbonates in Bermuda extend back to at least 880 ka and the subsurface units probably extend back to at least the Pliocene (Hearty et al. 1992; Vacher et al. 1995). Thus, for at least 2 million years, the only source of new sediment was carbonate precipitation, and its surface stratigraphic sequence comprises thick, cemented eolian carbonate dunes interbedded by thin paleosols (Vacher 1973; Hearty 2002). These two depositional modes are directly linked to phases of the Quaternary sea- level cycle (Sayles 1931; Harmon et al. 1978). As ice sheets melted five different times over the last half million years, sea levels rose, and Bermuda’s seamount was flooded to create dominantly marine system of shallow lagoons interspersed with small islands like the one we see today (**Fig. 1B**). During those high stand phases, carbonate muds accumulated in the lagoons from precipitation by algae, foraminifera, bivalves, and corals (Neumann 1965; Stanley and Swift 1968). At those times, the areal extent of terrestrial snail habitats was minimal and subdivided into small islands formed by the taller cemented dunes. As seas fell in at the onset of each new glacial phase, the unconsolidated carbonates were exposed, blown into dunes, and quickly cemented into massive eolianites (**Fig. 1D**; Vacher 1973). When sea level dropped below

**Figure 1.**
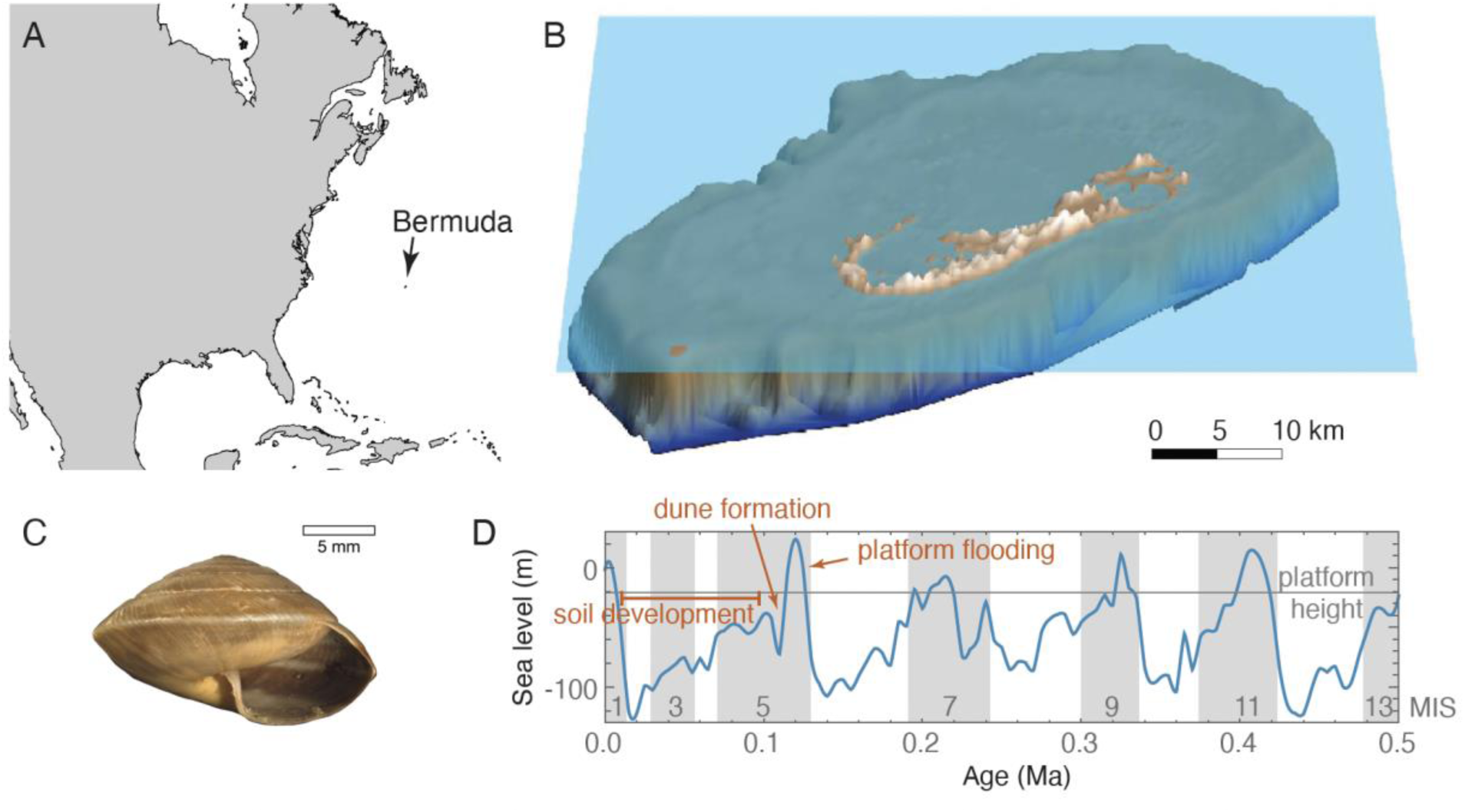
Overview. Map showing location of Bermuda (**A**). Rendering of Bermuda digital elevation model (DEM; Sutherland et al. 2013) with sea level set at approximately current height (**B**). Exemplar of a *Poecilozonites* snail shell (YPM IZ 104396, extant *P. bermudensis*) (**C**). Sea level for the last 50 kyr from Miller et al. 2005 showing the approximate height of the edge of the Bermuda platform (**D**).

-25m the entire Bermuda seamount top was exposed, forming a single large island without the shallow lagoons that characterize it today (**Fig. S1B**). The carbonate muds having already been dispersed or cemented, transportable sediment was unavailable during these glacial low stands, and the primary deposition was in the form of *terra rosas* soils a few centimeters thick in most places and no more than four meters thick in sinkholes or other karst depressions (Sayles 1931; Ruhe et al. 1961). At these times, the areal extent of terrestrial snail habitats was as much as an order of magnitude greater than at high stands. Each of these glacial low stand phases lasted about 100,000 years, about an order of magnitude longer than the phase in which water covered the seamount. Bermuda’s stratigraphic sequence thus consists of thick eolianites representing only hundreds or thousands of years of accumulation during the early falling stage of each sea level cycle, interspersed with thin paleosols, each of which may have taken as much as 100,000 years to form during the later falling stage, low stand, and early rising stage of each cycle (**Fig. 1D**).

To show how these evolutionary and stratigraphic processes might interact, I simulated a virtual metapopulation of *Poecilozonites* as it might have evolved by genetic drift through five glacial-interglacial sea-level cycles. *Poecilozonites* is assumed to have reached Bermuda during a single colonization event about 1 Ma (Pilsbry 1924; Hearty and Olson 2010). Three species groups, each of which was represented by one historically extant species and collectively comprised as many as 31 named fossil subspecies, diversified from the founder: *P. bermudensis* (**Fig. 1C**), *P. circumfirmatus*, and *P. reinianus* (Gould 1969). Today, *Poecilozonites* is critically endangered. *P. reinianus* disappeared by the 1950s and the remaining two species are nearly extinct in the wild (Bieler and Slapcinsky 2000; Outerbridge 2015). Based on its rich fossil record, Gould hypothesized that *Poecilozonites* had undergone many rapid speciation events as small peripheral populations were isolated. Bursts of morphological differentiation occurred in each founder due to drift and selection, after which the new species became established and underwent little morphological change. This model was the basis for his contribution to the punctuated equilibria theory of evolution (Gould 1969; Eldredge and Gould 1972). Some subsequent researchers have reinterpreted the fossil record of *Poecilozonites* as a single, anagenetic species that evolved by intense natural selection that arose from changing environments and turnovers in the types of predators that visited Bermuda (Hearty and Olson 2010). The present paper does not attempt to resolve how *Poecilozonites* evolved, rather it aims to use Bermuda and its snails to illustrate how sedimentary and microevolutionary processes can be controlled by the same causal factors, and how the correlation might make it difficult to disentangle rates of evolution and sedimentation. However, as discussed later in the paper, the outcome of this experiment offers a new perspective for interpreting punctuated patterns of evolution.

Using computational models, I show: (1) how the rise and fall of sea level across the platform’s edge restructures the spatial structure of populations on the islands through geographic range expansion, contraction, fragmentation, and coalescence; (2) how those spatial changes affect the outcomes of genetic drift via gene flow, founder effects, and genetic surfing; (3) how those metapopulation dynamics affect the morphological disparity of local populations and the rate and mode of evolution of the species as a whole; and (4) how the temporal and spatial processes of evolution correlate with an idealized sequence of sedimentary deposition. Computational modeling allows evolutionary parameters to be controlled that would be unknowable from the fossil record and allows the random evolutionary processes of interest to be separated from the confounding factors like selection and intraspecific competition that would be present in real evolving clades. Even though this model is based on the comparatively simple sedimentary and geographic history of Bermuda, I will argue that the results are generalizable to other situations, time periods, and drivers. I will also make the point that currently available statistical models of evolution are based on assumptions that do not explicitly account for the spatial processes that are the focus of this paper.

## Materials and Methods

### Computational Model

The computational model simulates the behavior of local populations of a single interbreeding species of snail in response to sea-level changes. The local populations can disperse, evolve (by genetic drift), interbreed with neighboring populations, and become locally extirpated. Each model runs from 0.5 million years ago to the present in 5,000 steps, each representing 100 years. During this interval there were five interglacials, including the present, when sea level flooded the Bermuda platform. When sea level is low, snail populations can expand across the entire seamount, but they are extirpated from flooded grid cells during late rising sea level phases (**Fig. 2A**). At each step of the model, every local population undergoes genetic drift, has a chance for offspring to disperse into an adjacent grid cell, interbreed with snails that disperse into its own cell, and experiences the possibility of extirpation, the probability of which increases if the cell is flooded (**Fig. 2B,C**).

**Figure 2.**
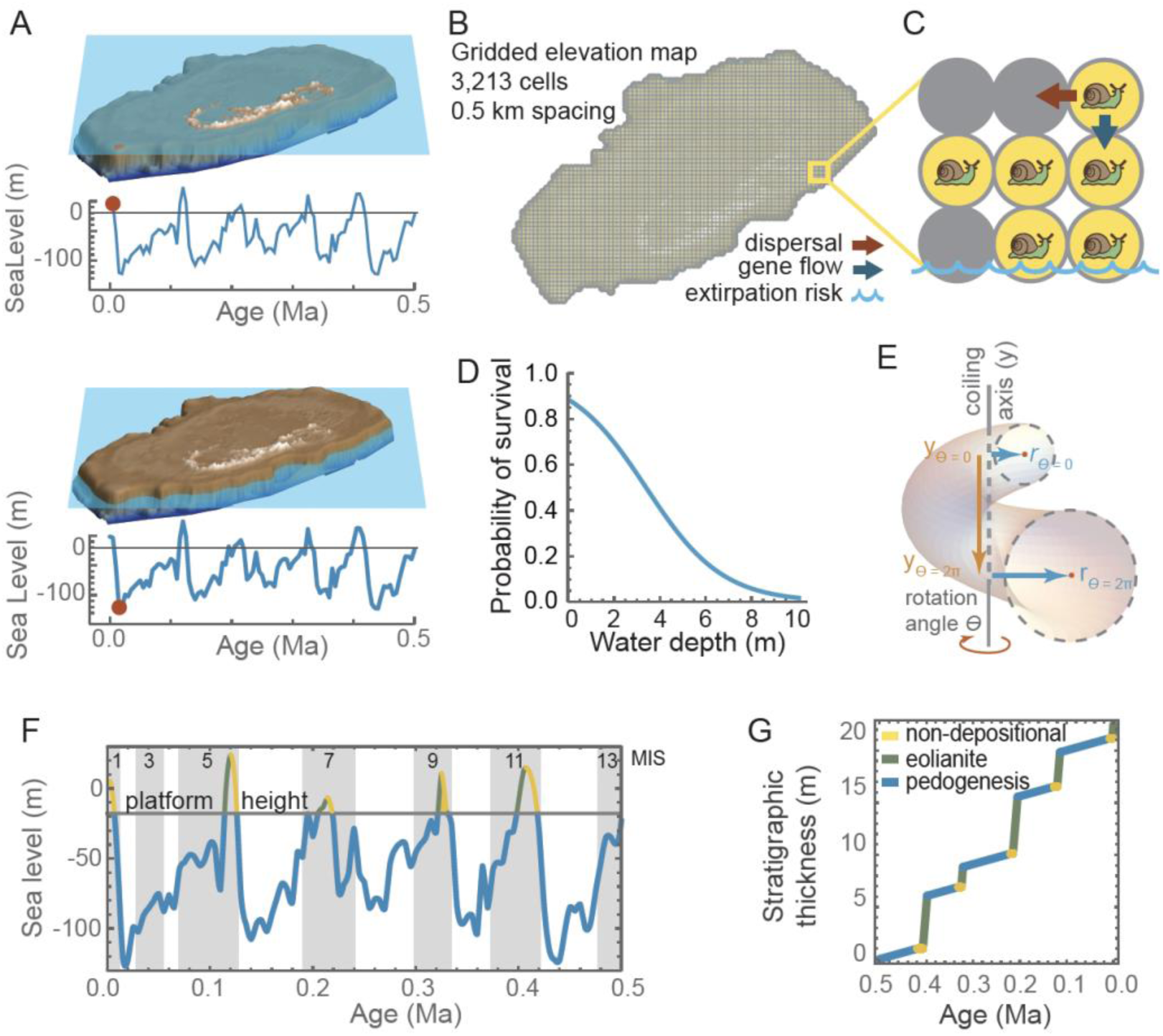
Computational model overview. The model simulates the rise and fall of sea level (**A**), the horizontal line approximating the height at which the seamount floods. The DEM is gridded into cells (**B**) that can be occupied by snail populations during dispersal events, they share morphologies through gene flow, and they become extirpated when a cell floods (**C**). Survival probability for a local population is 0.9 in fully terrestrial and declines to near 0.0 as water depth increases to 10 m (**D**). Snail morphology is modeled with Raup’s coiling equations (**E**). Time is classified into non-depositional (yellow), eolianite (green), and pedogenic (blue) phases based on the dominant sedimentary mode associated with phases in the sea level cycle (**F**). A sediment accumulation model was mapped onto time based on rates estimated from the thickness of Bermuda’s stratigraphic units (**G**).

A virtual landscape was created by gridding a digital elevation model of the modern Bermuda seamount (Sutherland et al., 2013) into a 100 x 69 cells, which are approximately 0.5 km per side or 0.25 km^2^ in area (**Supplement 1, Fig. S1**). This spacing is intentionally larger than an individual snail’s home range, but small enough that colonization of adjacent cells would easily occur within the century represented by each model step (cf., Baur and Baur 1993). Snails occupying each cell are treated as a single local population.

Sea-level change was modelled from the eustatic curve of Miller et al. (2005). Those authors estimated sea level for last 7 Ma at intervals of 5,000 years. I interpolated their data with a third order polynomial to model sea level at any point in time. The function was used to estimate water depth in each grid cell for each step of the model run. Marine isotope stage (MIS) boundaries follow Lisiecki and Raymo (2005). At today’s comparatively high sea level there are 165 terrestrial grid cells on the DEM model and only 44 at the even higher MIS 5e high stand (+44 m, 120 ka), whereas there are as many as 3,212 at the level of the MIS 12 low stand (–122 m, 435 ka). Anytime sea level is lower than –25 m in the model, more than 3,100 cells are available for snail populations.

Survival of local populations depends on whether their grid cell is flooded. When water depth in a cell is 0, probability of survivorship was 1.0. When water floods a cell, survivorship is scaled between 0.0 and 0.9 based on depth using the function *tanh*(-0.3*d* +1) (2+0.5)^-1^, where *d* is water depth in meters and *tanh* is the hyperbolic tangent function (**Fig. 2D**). The probability represents the possibility of unflooded points within the cell that might continue to support a snail population: survivorship is 75% or higher if water depth in the cell is less than 1.5 m (because the landscape is likely to still have many protruding islets), but nearly 0% when water is 10 m deep (because few cells would have subaerial habitat at that depth). During high stands snails can only persist on the highest topographic areas of the platform, during low stands water retreats from cells allowing snail populations to become established across the entire platform.

Shell morphology was modeled using five continuous-trait shell coiling parameters (*W*, *D*, *T*, *S1*, and *S2*), the meanings of which are described in detail below. At each step of the computation model, each of these traits evolved by drift in each local population. As described above, the rate of drift is *GN ^-1^*, where *G* can be rewritten as *h^2^σ^2^* (heritability × phenotypic variance, noting that *σ^2^* used here for phenotypic variance is a different parameter than the *σ^2^* rate of evolution discussed below). At each computation model step, each trait in each local population consequently evolved by a random amount drawn from a normal distribution with mean of zero and variance of *h^2^σ^2^N^-1^*. For all local populations, *N* was assumed to be 1,000 and *h^2^*was 0.5, both of which are biologically realistic values (Cheverud and Buikstra 1982; Polly et al. 2016). Values for *σ^2^*were chosen to allow the full range of known *Poecilozonites* morphologies to emerge over the course of a model run. As explained below, the step size was set to 100 years and the rate parameter scaled accordingly. To simplify the computations, *GN^-1^* was arbitrarily set to 1.0. and the variance of realized traits from each model were rescaled *post hoc* to the variance observed among the living and fossil Bermuda snails (see **Supplement 1**, **Table 1**). The resulting rates are equivalent to having using phenotypic variances *σ^2^* of *W* = 0.05, *T* = 0.05, *D* = 0.002, *S1* = 0.0004, *S2* = 0.0002 in combination with *N* = 1,000 and *h^2^* = 0.5.

**Table 1.** Summary of evolutionary model fitting. Mean AICC weight for each of the 12 evolutionary models across all five traits and all ten simulations is reported. *σ^2^* is the rate of evolution, *μ* is the directional parameter, and *K* is the number of parameters in the model, GRW stands for generalized random walk (i.e., a directional process), URW is unbiased random walk (i.e., Brownian motion), “same” means that the parameter was identical in the non-depositional, eolianite, and pedogenic phases, “all diff” means that the parameter was different in each of those phases, and “high diff” means that the parameter was the same in the non-depositional and eolianite phases, but different in the pedogenic phase. Models are sorted in order of their average support across all the simulations and traits.

| Test | Mode | $\mu$ | $\sigma^2$ | K | Mean Akaike Weight | Num Supporting Tests |
| --- | --- | --- | --- | --- | --- | --- |
| 1 | GRW | all diff | all diff | 6 | 0.05 | 0 |
| 2 | GRW | high diff | all diff | 5 | 0.14 | 0 |
| 3 | GRW | all diff | high diff | 5 | 0.01 | 0 |
| **4** | **GRW** | **same** | **all diff** | **4** | **0.35** | **23** |
| 5 | GRW | high diff | high diff | 4 | 0.03 | 0 |
| 6 | GRW | all diff | same | 4 | 0.00 | 0 |
| 7 | GRW | same | high diff | 3 | 0.06 | 2 |
| 8 | GRW | same | same | 2 | 0.00 | 0 |
| 9 | GRW | high diff | same | 2 | 0.00 | 0 |
| **10** | **URW** | **--** | **all diff** | **3** | **0.29** | **20** |
| 11 | URW | -- | high diff | 2 | 0.06 | 5 |
| 12 | URW | -- | same | 1 | 0.00 | 0 |
| 13 | URW/Stasis | -- | same/stasis | 3 | 0.00 | 0 |
| 14 | URW/Stasis | -- | all diff/stasis | 4 | 0.00 | 0 |
| 15 | GRW/Stasis | same/stasis | all diff/stasis | 6 | 0.00 | 0 |

Each local population disperses offspring into adjacent cells with a probability of 0.5 per cell per model step. Offspring carry their parent population’s parameters with them into empty cells but then evolve independently in subsequent model steps. If an offspring population enters an already occupied cell, it interbreeds with the established population and their trait values are averaged in the hybrid population to simulate gene flow. Each hybrid population is also assumed to have *N* = 1,000.

Each model run began 0.5 Ma (MIS 11) when sea level was below the platform (cf., **Fig. 2A**), thus allowing snails to become established across the island before the rising sea level flooded their habitats. At the beginning of each run, a single founder population was placed in a random grid cell with elevation of 5 m or higher with its trait values set to *W* = 1.7, *T* = 1.2, *D* = 0.05, *S1* = 0, *S2* = 0 (the latter being the average aperture shape for *Poecilozonites*). The run progressed through 5,000 steps to the present making the model step length one century long.

The modeling strategy follows Polly et al. (2016) and Polly (2019a). Code was written in *Mathematica* (Wolfram 2019) and is provided in both executable *Mathematica* notebook format and in readable PDF and text formats in **Supplement 2.** Simulations were run on Indiana University’s Karst high performance computing system with one CPU. Typical runs required 2.5 hours of computation, with additional time for output processing. Model runs were each assigned a unique name at runtime that begins with “Paleobiology” followed by the date and time and ending with a unique five letter random hash. Individual model runs are referred to by just the hash for brevity (e.g. SCRNT). The complete output of the models is available in **Supplement 3.**

### Snail Shell Modeling

The five traits are parameters for Raup’s (1966, 1967) shell coiling equations: *W* is whorl expansion, *D* is distance between coiling axis and aperture, *T* is the rate of translation along the coiling axis, and *S_1_*, and *S_2_* are geometric morphometric shape variables that describe the shape of the aperture (**Fig. 2E**). Cylindrical coordinates (*r* and y) for the shell at rotation angle are:

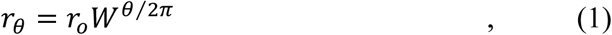

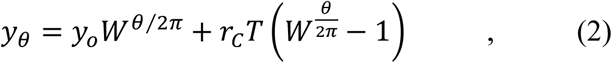

where *r_o_* is the distance of an aperture point from the coiling axis before rotation and *r_θ_* is its distance after a rotation to *θ*, *yo* is the original position of aperture points along the coiling axis and *y_θ_* is their position after rotation, and *r_c_* is the distance of the geometric center of the unrotated aperture from the coiling axis. The *D* and *S* terms are used to generate the *r*,*y* coordinates of the aperture so they do not appear explicitly in the equations.

Raup used coordinates of a circle for *S*, but any aperture shape can be used. I used the mean aperture shape *Poecilozonites*. As described in **Supplement 1**, I digitized the apertures of 35 living and fossil forms of *Poecilozonites* that collectively sample its full range of shape variation. Using geometric morphometrics, I derived two shape variables, *S1* and *S2*, from the first two principal components of the aperture morphospace (Bookstein 1991; Dryden and Mardia 1998). The mean shape is at the origin of any geometric morphometric morphospace, so the initial aperture shape was set at *S1* = *S2* = 0 for all model runs. Shape analysis was performed with *Morphometrics for Mathematica* v. 12.5 (Polly 2024).

Snail shells generated by the computational model were rendered by modeling the aperture from the *S_1_* and *S_2_* parameters by multiply them by the respective eigenvector and adding the consensus shape (see Polly and Motz 2017), converting to cylindrical coordinates, and offsetting by *D*. The aperture points were then plugged into Equations 1 and 2 with the simulated *W* and *T* parameters. Renderings were generated with the *Snails for Mathematica* v. 1.0 package (Polly 2022).

This simulation treated the shell coiling parameters as independent traits (e.g., as if they represent separate developmental genetic controls on shell morphogenesis), which means that they are fully independent of each other in the model output. In real snails, however, these particular parameters are not expected to be independent; they produce correlated effects on shell morphology that are difficult to disentangle if one were to back-estimate the same parameters from shells, and even in this simulation one would have to adjust for correlated traits if the phenotypic outcomes were assessed from the geometry of the simulated shells rather than from the parameter output themselves. To conceptually translate the patterns produced by these simulations to the real world, one would either need to choose traits that are known to be independent or use a phylogenetic comparative method that accounts for trait correlations (e.g., Revell and Harmon 2008).

### Output Processing

Raw model output was processed into three types of summary output. Animated maps were produced for each trait to show the distribution and trait values of local populations at each model step. Summary tables for each run contain the model time (age in millions of years), the number of extant local populations, a timestamp, and the mean, min, max, and variance of each trait across all the extant local populations. When there were fewer than five extant populations, the trait variances were truncated to 0.0. Finally, a series of graphs show the trait means and total morphological disparity through time. The results for all ten simulations are packaged together in **Supplement 3**.

### Geographic Variation Through Time: Standing Disparity

Standing morphological disparity among local populations was used as an index for geographic differentiation at each model step. Following Foote (1997), disparity was calculated as the summed variances of the populations across the five traits. Note that disparity of the shells in the mathematical space defined by the coiling parameters is not identical to the disparity of the rendered shells in a geometric morphometric space because the scaling between them is logarithmic and because the coiling parameters have interactive effects on the shell shape (see examples in Polly and Motz 2017; Polly 2017, 2023c). Regardless of how it is measured, however, the peaks and troughs of disparity would coincide despite a non-linear scaling in magnitude.

### Evolutionary Rates and Model Fitting

Rates and modes of evolution for the species as a whole were estimated using a modified version of the statistical evolutionary model fitting approach proposed by Hunt (2006, 2007). Hunt’s model fitting approach was applied to the data generated by the computational model to illustrate how the outcome might be interpreted if we encountered it in the fossil record. To apply Hunt’s or most other phylogenetic comparative methods, geographic and local variation in traits at any given time slice must be summarized as a mean and variance. The resulting time series of trait means represents an idealized pattern of the evolutionary behavior of the species over time, but ignores the spatial processes that produce the punctuated pattern of change. This tension will be discussed later in the paper.

The goal of the statistical model fitting was to determine whether the overall pattern of evolution fits a Brownian motion process (which one might expect given that the computational model uses a pure Brownian motion process to simulate genetic drift), directional evolution, or stasis, and whether it can be characterized as a single-rate process (which one might expect since the rate parameters for drift are held constant) or multi-rate process. Hunt’s approach accomplishes this goal by comparing the fit of three alternative evolutionary models to trait data sampled from an unbranching lineage: an unbiased random walk (URW) that is pure Brownian motion, a general random walk (GRW) which has a directional component, and a stasis model (stasis). URW is a typical Brownian motion model with one rate parameter (*σ^2^*). GRW is a directional model with two parameters, rate (*σ^2^*) and direction (*μ*). The stasis model is a type of adaptive peak or Ornstein-Uhlenbeck (OU) model with three parameters, rate (*σ^2^*), location (*θ*), and strength (*ω*). Note that Hunt’s *ω* is analogous to the *α* of many OU authors (e.g., Butler and King 2004), but Hunt’s model assumes that the lineage has already reached the adaptive peak and his *ω* parameter therefore represents only the stationary variance around the peak that emerges as a function of *σ^2^* and *a* in OU implementations like Butler and King’s. Here I refer to Brownian motion, directional evolution, and stasis as evolutionary “modes”. Likelihood is used to find the parameters for each model that best fit the trait data. The best model for each trait was selected using the sample-adjusted Akaike information criterion (AICC) and standardizing it into Akaike weights (Hunt 2006). Akaike weights sum to 1.0 and can be interpreted as the proportional support for each model.

I modified Hunt’s approach to test for differences in evolutionary rate and mode between the phases of the snail populations in which they are being extirpated from most of the platform during the late rising phase of sea level, when they are isolated in and then expand from high stand refugia during sea level high stand and early falling phase, and then when they cover the entire island during late falling phase, low stand, and early rising phase(**Fig. 2F**). Different rates and modes of trait evolution are expected during different eustatic phases because of the changing balance between local drift, gene flow, population isolation, and founder effects. Low stand phases were defined as the interval when sea level was below the edge of the seamount platform (-18 m). Bermuda was one large island during low stands (this is the phase of pedogenesis in the sedimentary cycle). Flooding phases were defined as the interval between when sea level surpasses -18m and the next high stand. During flooding phases snail populations become extirpated from low lying areas and persist isolated on the 10 to 15 small islands that remain above the high-stand sea level (this is also the phase of non-deposition in the sedimentary cycle). Regressive phases were the interval between high stand and when sea level falls below the platform edge. During the high stands there are fewer local populations, and the species is subdivided into isolated groups, potentially changing the population dynamic, compared to the low stands when snails expand across the island with gene flow across a very large number of local populations. During the regressive phases snail populations expand outward from high- stand refugia until they fill the entire platform with a single panmictic metapopulation (this is the eolianite phase of the sedimentary cycle).

For each model step, the mean value of each of the five traits was calculated across all populations extant at that time, yielding a 5000-step time series that traces the species’ overall evolutionary trajectory. Each trait lineage was binned into pedogenic (= late falling phase, low stand, and early rising phase), non-depositional (= late rising phase), and eolianite phases (= high stand and early falling phase. A series of “complex models” (Hunt et al. 2015) were then fit to each trait determine whether the rates or modes of evolution differed between eustatic phases. A total of 15 combinations of rate and mode were considered: (1) Generalized random walk (GRW; directional evolution) in which *σ^2^* and *μ* were different in each of the three phases; (2) GRW in which *σ^2^* was different in each phase and *μ* was different in high phases (non-depositional and eolianite); (3) GRW in which *μ* was different in each phase and *σ^2^* was different only in non- depositional and eolianite phases; (4) GRW in which *μ* was the same in all phases and *σ^2^* was different in each phase; (5) GRW in which *μ* and *σ^2^* were different only in non-depositional and eolianite phases; (6) GRW in which *μ* was different in each phase and *σ^2^* was the same in all phases; (7) GRW in which *μ* was the same in all phases and *σ^2^* was different only in non- depositional and eolianite phases; (8) GRW in which *μ* was the same in all phases and *σ^2^* was different only in the non-depositional and eolianite phases; (9) GRW in which *μ* and *σ^2^* were the same in all phases; (10) unbiased random walk (URW; Brownian motion) in which *σ^2^* was different in each phase; (11) URW in which *σ^2^* was different in non-depositional and eolianite phases; and (12) URW in which *σ^2^* was the same in all phases; (13) URW with different *σ^2^* in non-depositional and eolianite stages and stasis in pedogenic phase; (14) URW with the same *σ^2^* in non-depositional and eolianite stages and stasis in pedogenic phase; (15) GRW with the same *µ* but different *σ^2^* in non-depositional and eolianite stages and stasis in pedogenic phase (see also **Table 1**).

Evolutionary model fitting was carried out in *Mathematica*© using functions available in *Phylogenetics for Mathematica* v. 6.8 (Polly 2023a). In its distributed form, the *ThreeModelTest* function does not fit complex models so analyses were customized using its sub-functions. The complete code found in **Supplement 2.**

As will be discussed later in the paper, there is a tension between the spatial processes used in my computational model and the assumptions that underpin evolutionary statistical models, including the one used here and most if not all standard phylogenetic comparative statistical models. The statistical models implicitly assume that each lineage in a data set or on a phylogenetic tree behaves like a single population at each time step, and the evolutionary rate estimated from fitting the model to real data is based on that assumption. My computational model involves many local populations, each with a constant rate of evolution, that interact at each step. As discussed below, the rates of evolution for the species as a whole as estimated by fitting Hunt’s statistical evolutionary model will be different from the rate in the local populations.

### Comparison of Species-level Evolution to an Idealized Stratigraphic Column

Sedimentation was not included in the computational model, but the results of the model were compared to an idealized stratigraphic column constructed from the modes of sedimentation that dominantly occur on Bermuda during each phase of the sea level cycle: high stand = carbonate production; early falling phase = eolian dune formation; late falling phase, low stand, and early rising phase = soil formation; late rising phase = no sedimentation. Just like the average trait values that were used for evolutionary model fitting, this idealized column represents what we might expect the local stratigraphic thicknesses to be like at a hypothetical location where fossils might be sampled. Bermuda’s stratigraphy is spatially complex because it is dominated by eolianites of heterogenous thicknesses that have not only vertical but lateral superpositional relationships and soils that discontinuously blanket low-lying areas, the topography of which is also affected by karstification (Vacher et al. 1995; Hearty 2002). In real situations, sedimentary accumulation will vary locally in ways that will complicate the relationship between sedimentary and evolutionary rates.

The intention of this paper, however, is simply to demonstrate in a theoretical sense how a single physical factor like sea level can exert a correlated effect on both microevolutionary processes and mode of sedimentary deposition. To do this, I correlated the computational model’s absolute ages to the generalized stratigraphic column so that evolutionary changes could be plotted as they might be inferred by a paleontologist using a stratigraphic meter-level system (**Fig. 2G**). I created the idealized stratigraphic column by setting its total thickness to 20 m (approximately the median total thickness of Bermuda’s surface units) and subdivided it into eolian, pedogenesis, and non-depositional phases whose temporal lengths were derived from the sea-level curve (**Fig. 2G**). The rates in each phase were based loosely on the observed stratigraphic thicknesses of Bermudian units and the lengths of time over which they accumulated (Vacher et al. 1995; Hearty 2002). The eolian phase occurs during early falling sea level phase when rapid accumulation of wind-blown carbonate sands and muds were cemented into thick eolianites, the rate of which was based on the Southampton Fm. that accumulated as much as 10 m thickness in less than 10,000 years during MIS 5a and was thus modeled at 0.001 m yr^-1^. Low stand pedogenesis was modeled at 0.0005 m yr^-1^ based on the estimated 0.5 m thickness of the St. George’s geosol, which accumulated over approximately the 100,000 years of MIS 2-4. The non-depositional phase occurred in the late rising phase of sea level which rapidly flooded the platform, reworking some units into beach conglomerates but otherwise producing no accumulation, was modeled with a sedimentation rate of 0.0m yr^-1^. Pure carbonate deposition in the high stand lagoonal areas was not included in the idealized stratigraphic column because it only appears in transported form in the sections that are currently above sea level as eolianite.

The mean trait values from the evolutionary model fitting exercise were rescaled from their original time units to expected stratigraphic thickness based on these idealized rates of sedimentation. While the approach is simplistic, the interpretations I draw from it only hinge on the observation that the snail fossils found preserved in the thick eolianites of Bermuda accumulated rapidly in the short regressive phases and the fossils in the thin paleosols accumulated slowly over the long glacial low stands.

## Results

### Morphological Disparity Increases During High Stands

Two questions about trait evolution are of interest here: does standing morphological variation change as sea-level cycles progress, and how does the average morphology evolve through a series of cycles?

The first of these, morphological variation, manifests itself as geographic variation among local populations and can be measured as disparity. In the model results, geographic disparity is visibly linked with phases of the eustatic cycles, with disparity falling to almost zero during low stands and increasing during the short high stand and early falling phases (**Fig. 3**). Disparity here is calculated as the variance in a shell trait across all the local populations extant during each step of a model run. Disparity of one trait (*T*) during one model is illustrated in **Fig. 3A-F**, and disparity averaged across all traits and all model runs is shown in **Fig. 3G**.

**Figure 3.**
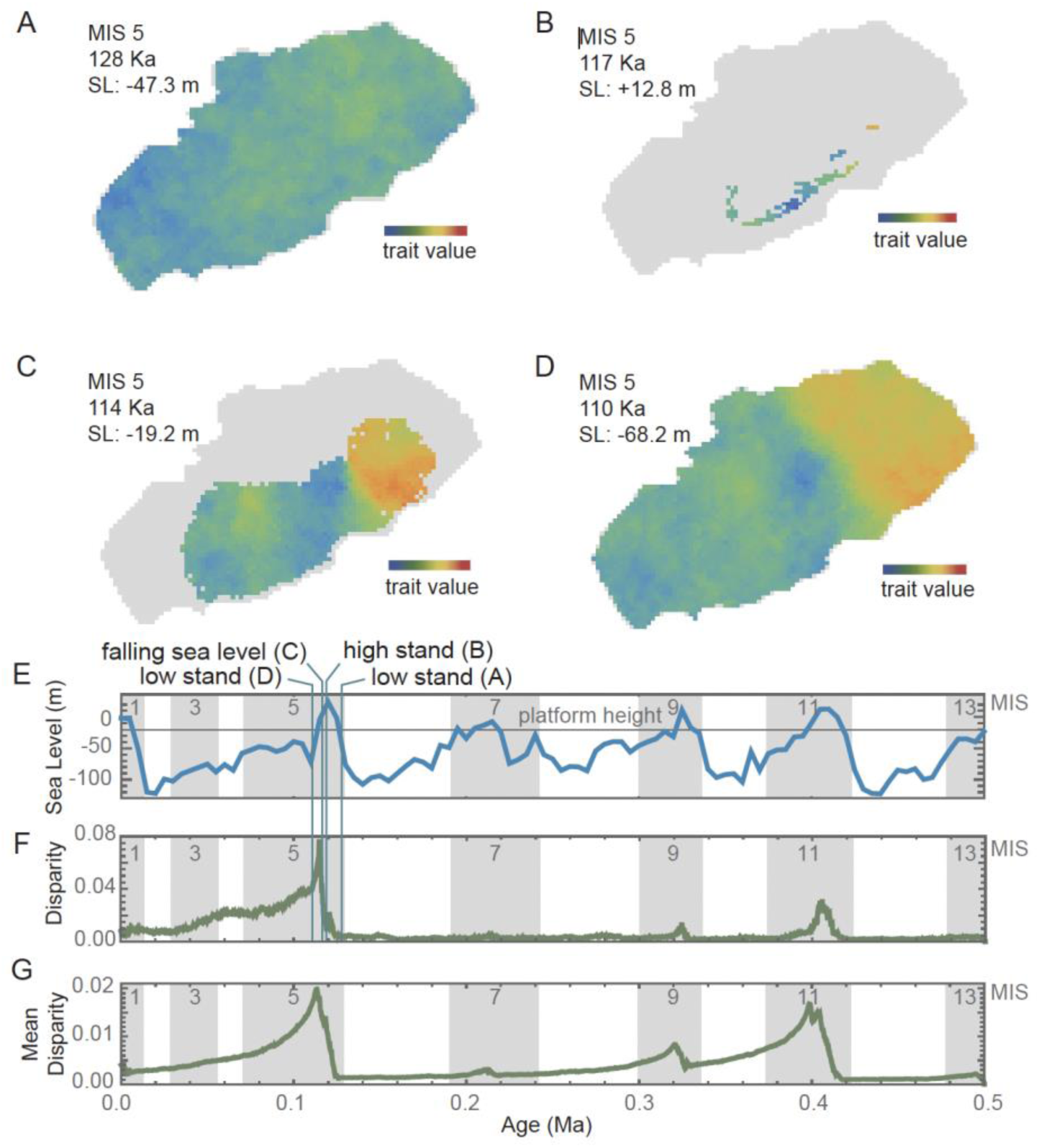
Geographic disparity through time. Snapshots of geographic disparity in a single trait through the phases of the glacial-interglacial eustatic cycle at low stand (**A**), high stand (**B**), population expansion associated with the regression phase (**C**), and the initial pattern at the beginning of the next low stand (**D**). Over the duration of the 0.5 Myr run, geographic disparity (**F**) rises at high stands (**E**) and slowly during low stands. The vertical lines in E and F show the time slices A-D, all of which are illustrated with the trait T from model run DOLTH (this trait and model run were chosen arbitrarily for illustration). All 5 traits and all 10 model runs produced similar disparity patterns as shown by their mean disparity (**G)**.

During low stands, disparity is low because as local populations occupy the entire platform their traits equilibrate through the homogenizing effects of gene flow (**Fig. 3A**). While disparity is low at those times (**Fig. 3E-G**), it never reaches zero (complete homogeneity) with the parameters used in the computational model because genetic drift in local populations adds white noise faster than gene flow can average it out (**Fig. 3A**). The minimum level of disparity during low stands is thus controlled by the balance of genetic drift, which causes each local population to become randomly different from its neighbors, and gene flow, which causes local populations to become more like their neighbors, something that found not only in this computational model but also in real populations (Levins 1968; Endler 1977; Polly and Wójcik 2019).

Disparity rises along with sea level (**Fig. 3E-G**). As seas flood the island at the end of each glacial period, it isolates small numbers of local populations on island refugia where their traits quickly diverge due to drift in the absence of gene flow (**Fig. 3B**). A short lag is observed between rising sea level and rising disparity because disparity starts increasing once islands become isolated and is then augmented through the edge effect of “genetic surfing” as populations expand from island refugia as sea level drops (**Fig. 3C**). Morphological clines that were seeded by “surfing” from the differentiated refugial populations maintain some morphological disparity that is steadily lost as gene flow re-homogenizes the populations, slowly returning the system to the same non-disparate state that preceded the interglacial phase (**Fig. 3D**).

Disparity at high stands and its rate of homogenization depend on the rates of drift and gene flow. I purposefully selected parameters that maximized the difference in disparity between high and low stands to illustrate the process. The same relationship would emerge no matter what parameter values were chosen but note that these parameters interact with cycle length in a complex way as discussed later in this paper.

### Trait Values Shift During High Stands

Trait means shift abruptly during high stands, in seeming contradiction to the Brownian motion nature of genetic drift and the constant rate of drift at the population level. The trait mean captures the central tendency of evolution around which geographic variation fluctuates, and it is the parameter from which evolutionary rates and modes are normally inferred (cf., Simpson 1944; Lande 1976; Felsenstein 1988; Gingerich 2001; Polly 2004; Hannisdal 2006; Hunt 2006; Walsh and Lynch 2018).

The pattern of trait evolution in these models (**Fig. 4**) is much like that expected from punctuated equilibria: long periods with little trait change punctuated by rapid bursts of change (cf., Eldredge and Gould 1972; Gould and Eldredge 1977). The difference is that speciation is absent in this computation model but integral to Eldredge and Gould’s. Like with trait disparity, the punctuated pattern of trait change in the computational model is linked to sea level. During low stands, traits evolve slowly in the large, panmictic metapopulation that covers the seamount, punctuated by rapid bursts of change in the small, isolated populations that survive during high stands. The bursts are most apparent at the MIS 11 and 5 high stands, and to a lesser extent at MIS 9. The rapid phases of trait evolution arise from new morphologies that appear on the isolated island refugia and are swept to dominance by “genetic surfing” by population expansion during the eustatic regression phases (**Fig. 3C,D**). Gene flow then homogenizes the species around a new trait mean during the low stands (**Fig. 4**).

**Figure 4.**
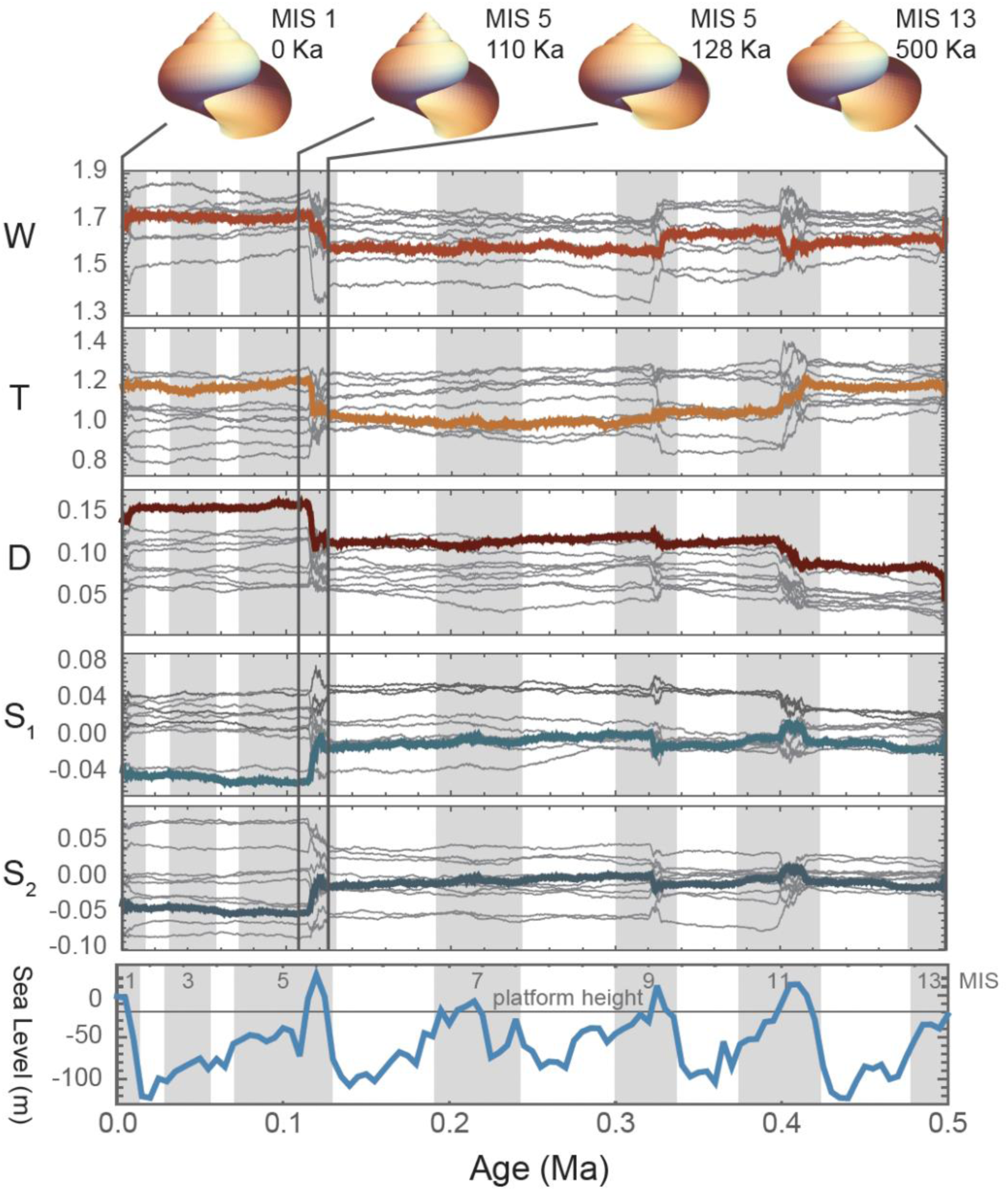
Trait change. Mean values of the five traits through time from model run DOLTH (thick colored lines) superimposed on mean values of the other nine runs with sea level for context. Each line represents the mean trait value across all extant populations at each model step. Four shape models are shown with vertical lines indicating their location in time (note that the 3^rd^ and 2^nd^ models correspond in time to **Figs. 3A and 3D** respectively).

Punctuated Patterns are Produced by Rate Shifts, not Changes in Mode Punctuated patterns of trait evolution occur at high stands at approximately the same time as the peaks in morphological disparity (**Fig. 4**). The “punctuation” events are produced by increases in evolutionary rate, not by episodes of relaxed stasis as shown by evolutionary model selection. The models that best fit the trait data were ones in which the evolutionary rate (*σ^2^*) differed between non-depositional, eolianite, and pedogenic phases (tests 4 & 10), which together accounted for 43 out of 50 of the traits simulated in this study (**Table 1**). In the remaining seven cases the evolutionary models that were supported were ones in which the non- depositional and eolianite phases (“high stand”) shared a common rate of evolution that differed from the low-stand phase (tests 7 & 11).

A summary of evolutionary rates (*σ^2^*) broken down by sea-level phase is found in **Table 2** (see also histogram summary in **Supplement 1, Fig. S4**). The two high-stand evolutionary rates were always at least one order of magnitude faster than the low-stand rates, and the regressive phase was usually about 1.5 times faster than the non-depositional phase. Note that by design the rates of the *D* and *W* traits were higher than *T*, *S1*, and *S2* traits to explore the full range of observed values for Bermudian snails in each run.

**Table 2.**
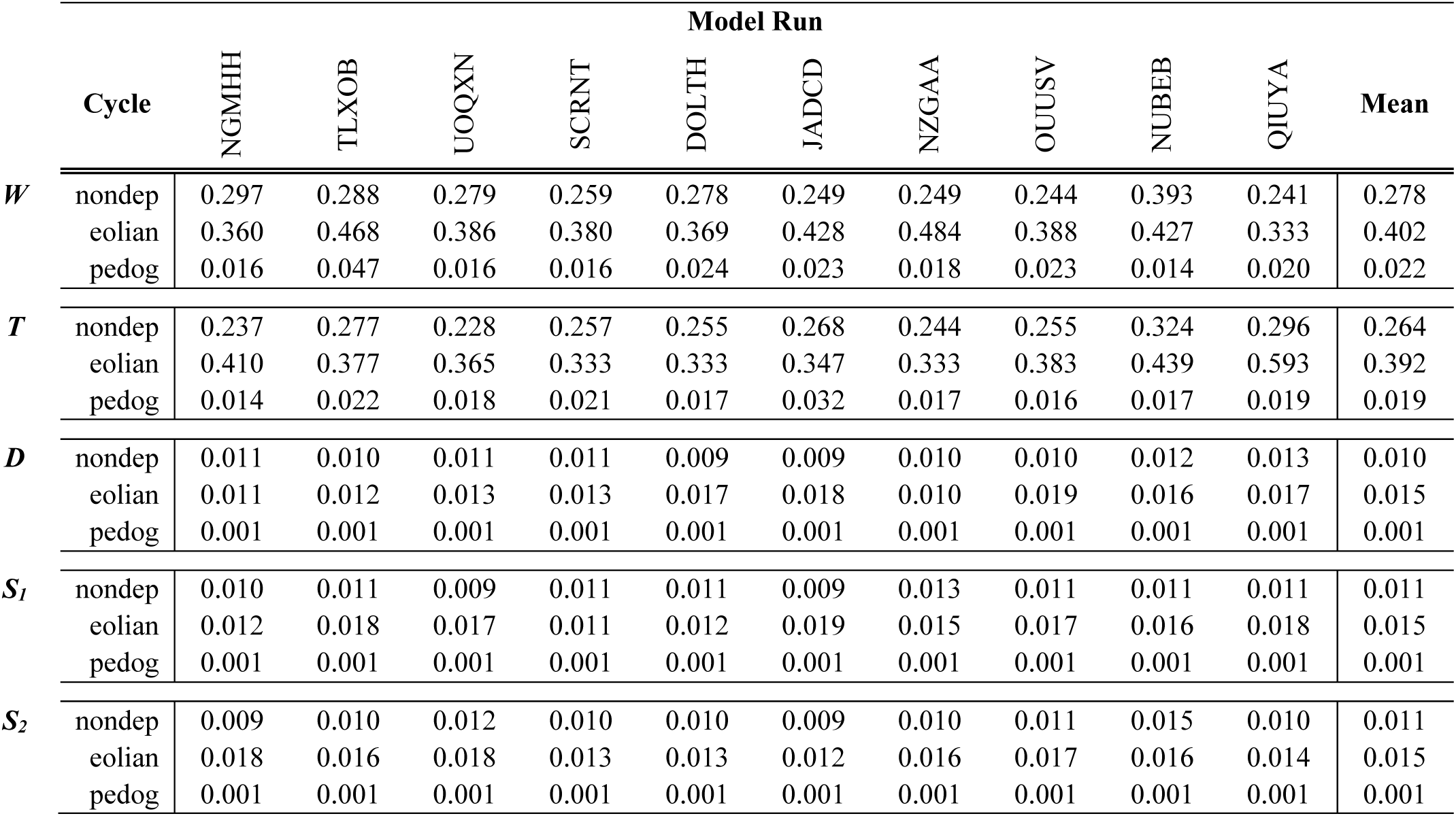
Evolutionary rate (*σ^2^*) results. For each of the five traits, the mean evolutionary rate and the rate for each model run are reported for each phase of the sedimentary cycle. The rate of trait change is at least one order of magnitude slower during pedogenic phases (when sea level is below platform height) than either non- depositional or eolian phases.

## Discussion

### How Does a Multiple-rate Pattern Emerge from a Single-rate Process?

Single-rate Brownian motion (BM) models (test 12) were never supported (**Table 1**) even though the underlying genetic drift process coded into the computation model is inherently a single-rate BM process in which all the local populations drift randomly at the same rate.

Instead, the rate of evolution that emerged from the computational models was always different between low stands and high stands (50 out of 50 times), and usually different between non- depositional, eolianite, and pedogenic phases (43 out of 50 times). In fact, evolution was almost always an order of magnitude faster during non-depositional and eolianite phases than in pedogenic phases.

The reason is that the rate of evolution of the species as a whole (i.e., the trait mean) is a function of the total number of interbreeding local populations and their geographic contiguity, both of which are affected by sea level. When there many populations, the overall or effective population size of the species as a whole (*Ne*) increases, which lowers the rate of drift (Whitlock and Barton 1997). The overall rate of trait change is slowed because the random changes in local populations are averaged out by gene flow from neighboring cells, which is a function of the number of occupied cells and the rate of dispersal (*Nm*). When only a small number of isolated local populations exist at high stands, they rapidly diverge because the rate of drift on each islet increases inversely to the number of local populations and because gene flow ceases between populations isolated on different islets (**Fig. 3B**). The newly acquired variety of trait values are swept into the species as a whole through “surfing” events as the islet populations expand over the seamount during regressive phases (**Fib. 3C,D**). Lowered *Ne* increases the rate of drift within each isolated local population cluster, the breakdown in gene flow allows disparity to increase, and surfing carries the high-stand differentiation into the enlarged metapopulation at the beginning of each low stand. Radically different rates of evolution thus characterize each eustatic phase.

The multi-rate behavior of evolution in this system is the result of what are fundamentally “Wrightian” spatial dynamics of a metapopulation (Polly 2019a). The standard evolutionary models like BM, OU, and directional selection, including those applied here (e.g., Paradis et al. 2004; Hunt 2007; Beaulieu et al. 2012; Clavel et al. 2015), are based on the statistical expectations of a single evolving population, the fundamentals of which are derived from Lande’s (1976) formulation for evolution of quantitative traits in response to selection. In its standard form, this “Fisherian” model describes the behavior of a single, unstructured panmictic population using the principles outlined in the classical quantitative derivations of Fisher (1930). Structured populations consisting of local populations, each with its own trait mean, that interact through dispersal and interbreeding, and whose number varies because of local extinction and colonization were considered by Wright (1931, 1940) to explain evolution in the natural world, concepts that later were elaborated into the field of metapopulation ecology (e.g. Levins 1968; Hanski 1999). While metapopulation processes are all “microevolutionary” in nature, the interaction of a metapopulation with changing environments over geological timescales can produce “macroevolutionary” patterns like the punctuated events driven by fragmentation of a species’ geographic range in these computational models. It seems likely that such spatial interactions with Earth system processes have been the norm in the history of life, which may mean that the current “Fisherian” evolutionary models do not adequately describe the patterns of trait evolution found in the fossil record (Polly 2019a, 2020). That the traits of local populations had to be averaged across the entire island to apply Hunt’s model fitting method and that it supported a multi-rate model of evolution in which rate changes are tied to expansions and contractions of the geographic range of the metapopulation rather than true changes in rate parameter used in the computation model, which was constant through time and across populations, illustrates this tension. Evolutionary model fitting is still useful, as is demonstrated by the application of Hunt’s method to the computation model data, but users might well consider a broad range of scenarios when interpreting results like the ones modeled here in which there is a constant microevolutionary rate at the local population level and an emergent shift in rate at the species level driven by change in geographic range size rather than change in selection intensity or mutation rate.

### Why is Slow Trait Change at Low Stands not “Stasis”?

The patterns of overall trait evolution produced by the computational model visually look like stasis, with long periods in which little or no change is punctuated by rapid bursts of change during high stands (**Fig. 4**). Eldredge and Gould (1972) did not originally define stasis, except as a history of very little change within a species relative to the degree of difference between that species and its ancestor at the time of speciation. In that general sense, the pattern emerging from this spatially explicit model of evolution is ostensibly one of stasis punctuated by rapid bursts of change.

Later stasis and directional evolution were statistically defined as accumulating either less or more differentiation than expected under a Brownian motion model (Lande 1986; Bookstein 1988; Gingerich 1993), the statis definition similar to the now-familiar adaptive peak (OU) evolutionary models. Hunt’s model selection algorithm shows that the slow low-stand phases are in fact Brownian motion but at a slow rate, with true stasis models never being supported out of 50 different trait runs (**Table 1**). The reason is that even with large numbers of populations that average their phenotypes by gene flow, the net effect of local drift processes produces Brownian motion in the species’ mean (Whitlock and Barton 1997). This appearance of stasis is an illusion of scale between the low-stand and high-stand rates, which differ by more than an order of magnitude (**Table 2**).

### Why do “Directional” Evolutionary Models Sometimes Fit the Computational Model Outcomes?

Directionality is completely lacking in the computational model, yet it sometimes emerges as a best fitting evolutionary model (GRW, tests 4 & 7). Directionality in these results is due to change in which the large evolutionary burst in MIS 5 creates an overall “directional” trend (e.g., **Fig. 4**). This interpretation is supported by the fact that only single-*μ* parameter GRW models fit the data (tests 4 & 7), never multiple-*μ* models. The Brownian motion component is almost always about an order of magnitude more important than the directional component. To see this, compare typical values of *μ* and *σ^2^* in **Supplement 1, Fig. S5** (note *μ* must be squared to make its units comparable to *σ^2^*). The directionality parameter *μ* is almost always near zero with its variance in the range of plus and minus 0.1 (squared = 0.01), whereas the evolutionary rates *σ^2^* in the high-stand phases are always greater than 0.2.

### Extending the Common-cause Hypothesis

The common-cause hypothesis posits that the same tectonically and climatically driven sea- level changes that control the structure of the rock record also control biodiversity patterns (e.g., Newell 1952; Sloss 1963; Hallam and Wignall 1999; Peters and Foote 2011; Peters and Heim 2012). One common cause factor is the degree of continental flooding. When a continental shelf floods at high stands, accommodation space for the deposition of new sediments increases along with the availability of shallow shelf habitats and opportunities for speciation, but when the shelf is exposed at low stands, deposition ceases over a wide area and shallow marine habitats are lost resulting in widespread extirpations and extinctions. The volume of the rock record is therefore correlated with standing diversity in this example. This type of common cause is hypothesized to be one of the primary drivers of macro-evolutionary processes that operate above the species level driving the rise and fall of entire clades, causing selective mass extinctions, and sorting taxa by shared functional traits (e.g., Jablonski 2000, 2017).

This paper presents an extension of the common-cause hypothesis, that the same geological controls that affect biodiversity by altering macroevolutionary rates of speciation and extinction may also affect processes of microevolution at population levels by altering the rate of evolution within species. The mechanism is similar – the impact of expanding and contracting habitats – but here the tempo (and perhaps mode) of evolution within evolving species lineages are hypothesized to be controlled by their effects on population size and thus the importance of drift and the rate at which it changes the phenotype of the species as a whole.

While natural selection has not been discussed in this paper, it is more effective in large populations because the drift component of evolution decreases and because the amount of genetic variation increases (e.g., Wood et al. 2016). Large populations are also subject to “selective sweeps” that spread neutral or deleterious alleles along with beneficial ones, thus rapidly changing the genetic nature of a species in unexpected ways (e.g., Stephan 2019). The concept presented in this paper thus probably applies to non-neutral microevolution as well as to evolution by pure drift, although in more complex and less predictable ways.

### How Might Evolutionary and Stratigraphic Processes be Conflated in the Fossil Record?

The primary message of this paper is that the same processes that control deposition in the sedimentary record may also control microevolutionary processes like the rate of evolution. In the Bermudian system, the expected rate of drift peaks during regressive phases, which is also precisely when the eolian phase of dune accumulation occurs, and conversely the rate of drift slows at the same time as sedimentation turns to pedogenesis during the low stands.

If trait change were to be read straight from the sedimentary record, how would the pattern be distorted by the interaction? **Figure 5** shows a side-by-side comparison of change in the mean value of the *T* parameter by scaled by time and by sediment thickness in the idealized stratigraphic column. Overall, the punctuated pattern of trait change is smoothed into a more gradual trend when read by meter level because the rapid regression bursts are stretched over thick rock units (green lines) and the long low-stand stretches of slow change are compressed into thin soils (blue lines). Punctuation is accentuated during late rising phase and high stand (yellow lines) when rates of evolution are high, but sediment accumulation is theoretically at its minimum. This system would therefore bias paleontological interpretations toward a gradual evolutionary pattern with some punctuated events but little evidence of stasis-like periods of little change. Gould (1969) was mindful of variable sedimentary rates, but if the sea level really did influence both evolutionary rate along with sedimentary deposition in the Bermudian record it would have made the evolutionary pattern appear more like gradualism rather than punctuation and stasis and thus his interpretation of punctuated equilibria is, in principle, robust relative to the potential conflation illustrated here.

**Figure 5.**
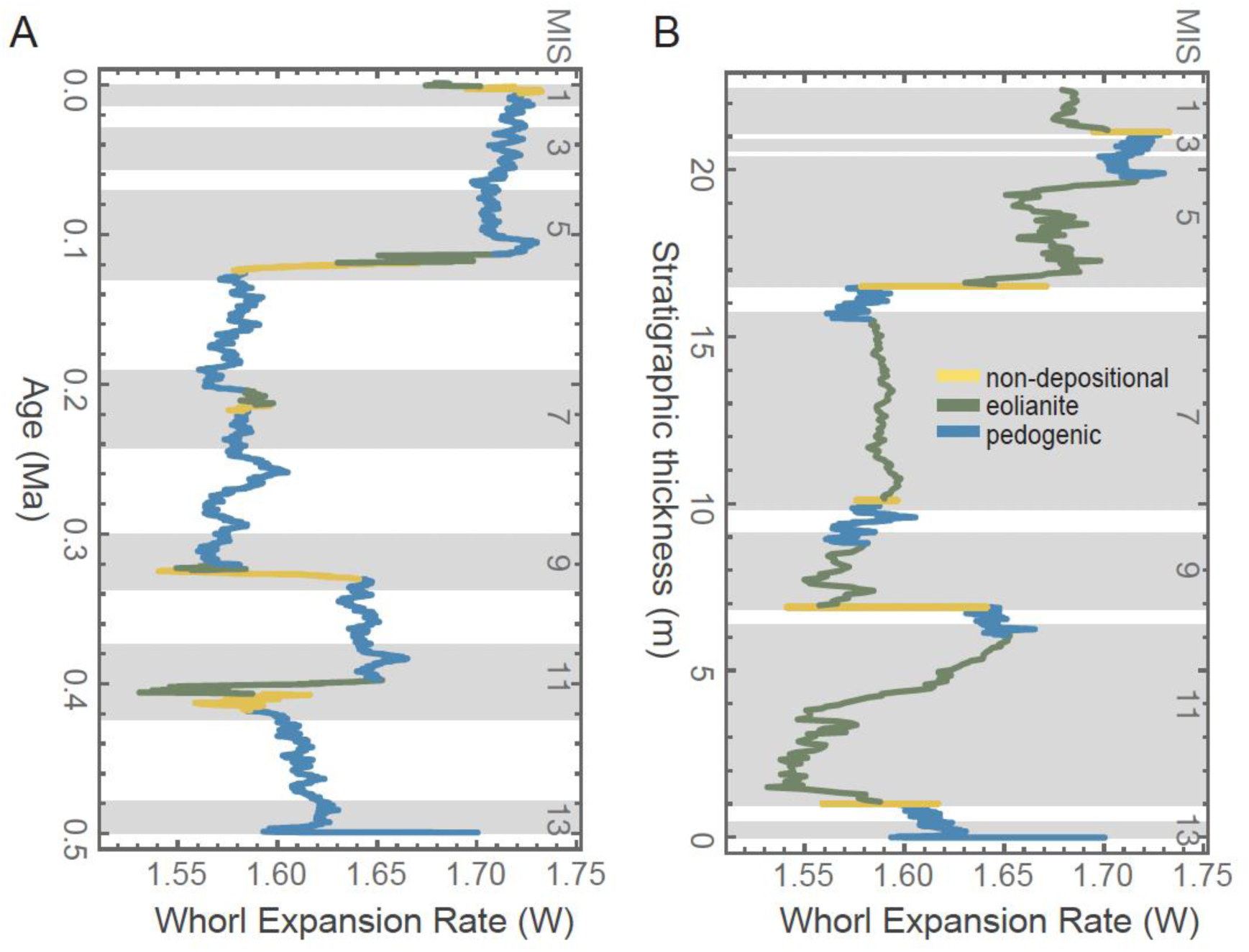
Temporal and stratigraphic scalings of trait change. Trait change scaled to real model time (trait T from model run DOLTH as in **Fig. 3**) shows strong punctuated rate changes at non-depositional (yellow) and pedogenic (**green**) stages (**A**), but the pattern is distorted into a seemingly more “random” model when scaled by an idealized stratigraphic thickness (**B**).

Paleontologists have, of course, been conscious that small population sizes may produce both higher rates of evolution and lower rates of preservation in the fossil record. Not only did that scenario feature strongly in Eldredge and Gould’s (1972) formulation of punctuated equilibrium, but so too in Simpson’s “quantum evolution” scenario, “In small populations undergoing pronounced shifts in environment and ecology, much higher rates of evolution are possible…. From their very structure, such groups do not leave good or continuous fossil records….” (Simpson 1944, p. 119). Less attention has been paid to whether the processes that might produce small populations or rapidly changing environments are causally linked to the processes that control sedimentation rates or preservation potential. Hunt (2008) pointed out that the density of sampling in the stratigraphic record can determine whether a rapid burst of evolution appears instantaneous (“unsampled punctuation”) or directional (“sampled punctuation”). The rapid bursts of trait change shown during rising sea level phases in **Figure 5,A** are good examples of what would be an unsampled punctuation in the fossil record since these rapid bursts of change take place during what would likely be a period of non-deposition in the Bermudian system. The rapid bursts of trait change during falling phases, however, would be examples of “oversampled punctuation” because what would have been a rapid event is drawn out across what would be the most productive depositional phases in the Bermudian sequence, the thick eolianites. The low stand pattern in which long periods of slower evolution are stratigraphically compressed into thin soil units could be termed “static compression”. The overall effect of translating the temporal pattern of changes in **Fig. 5,A** into expected stratigraphic thicknesses in **Fig. 5,B** is to dampen the strong punctuated pattern in the former into what appears to be a more of a “gradualist” random walk in the latter.

Sedimentologists and paleontologists have been conscious that sedimentation rates vary widely and sometimes undetectably through a section (e.g., Kraus and Gwinn 1997; Goddard and Carrapa 2018; Valenza et al. 2022). Hunt (2004) discussed its implications for measuring evolutionary rates in the fossil record. Recognizing that fossils must be binned at some level, Hunt showed that the time averaging imposed by analytical stratigraphic binning has the effect of increasing the appearance of within-bin variance and decreasing between-bin evolutionary change when the bin represents longer periods of time, like with the low-stand phases of this simulation. Here, however, the lengthy “bins” of the thin paleosol units is offset by their slower evolution.

### What Would the Disparity Pattern Look Like with Speciation?

This exercise does not directly test Gould’s (1969) hypothesis that Bermudian snails evolved under a punctuated equilibria model. While the modeling exercise suggests that the null expectation of random evolution on Bermuda might be long periods of very slow change punctuated by bursts of rapid evolution, punctuated equilibria is a hypothesis that evolution is rapid at speciation and speciation was intentionally omitted from my computational model in order to focus on how Earth system processes might impact “anagenetic” evolution within a single species lineage.

How would the model change if it involved speciation? The probability of speciation would also increase during high stands because populations diverge on small, isolated islets and thus are more likely to develop the barriers required for reproductive isolation. In the model, no such barriers are included so gene flow begins to erase the differentiation that accrued in isolation once the populations re-expand to cover the seamount at low stand. A peak in disparity is thus observed at high stands that decays over the course of the next low stand (**Fig. 3F,G**). If true speciation occurred during the high stands, we would expect disparity to grow just like in the model, but it would not be lost during the next low stand because the now disparate species would not coalesce through gene flow. The increased disparity would thus be retained until the next high stand, when isolation would once again occur, increasing disparity even further, and would once again be maintained during the low stand. The expected disparity pattern with speciation would therefore be a stair-step pattern in which disparity jumps up at each high stand (extinction and incomplete speciation would, of course, add complexity to the pattern).

Gould (1969) argued that a pattern like this is found in the Bermudian fossil record. In contrast, Hearty and Olson (2010) later argued based on new geochronological work that the disparate fossil forms of *Poecilozonites* were actually sequential in time rather than contemporaneous and that they represent a single anagenetic lineage similar to the ones in my computational models. While the empirical data in Bermuda’s fossil record could be used to test the punctuated equilibrium hypothesis using the two models of disparity with and without speciation, more work is required. The author notes, however, that two or more forms of snail sometimes occur cemented together in the same sinkhole breccia suggesting that they were both contemporaneous and coexisted in the same habitat, strongly implying speciation.

## Conclusions

What null model should we assume for trait evolution? Hypothesis testing in evolutionary morphology usually pits a hypothesis implying some causal process, like directed change or stasis, against a null model of “no process”. Since the ground-breaking work of Felsenstein (1988), Bookstein (1988), and Gingerich (1993), the null model for trait evolution has been a single-rate Brownian motion process, not because it is believed to be the dominant way in which evolution proceeds, but because it invokes no particular process and implies no directionality, no correlation, and a rate that varies stochastically around a single mean. This paper suggests that a variable rate Brownian motion null model might be more appropriate. As discussed at length elsewhere, the typical single-rate Brownian motion model when applied at phylogenetic time scales is inherently a “Fisherian” model that assumes that species behave as single, panmictic populations with a uniform phenotypic mean and variance and a stochastically constant population size (Polly 2019a, 2020). Real-world evolution is likely to be “Wrightian” in which species are metapopulations whose means vary spatially, whose ranges expand and contract, and whose population parameters – and therefore rates of evolution – change in response to changing environments. It should be realized, however, that the Wrightian model presented in this paper makes its own process-based assumptions, namely that true genetic drift is the dominant process in evolution. A fluctuating selection null model (Kimura 1954; Felsenstein 1988), for example, might behave more like the classic single-rate Brownian motion model embedded in most current phylogenetic comparative methods.

The hypothesis put forward in this paper is that the same causal processes that control sedimentary regimes, habitat availability, and climate configurations – whether those be Milankovitch cycles, tectonics, or patterns of oceanic and atmospheric circulation – not only exert control on standing biodiversity through their effects on rates of speciation and extinction, but that they may also affect rates and modes of trait evolution. If so, then the common-cause hypothesis may extend into the realm of microevolution (population-level processes) as well as macroevolution (species- and clade-sorting and ecosystem-level processes).

The mechanism by which Milankovitch-driven sea-level cycles would affect trait evolution in a terrestrial species like *Poecilozonites* and drive sedimentation patterns is an obvious and perhaps an extreme example. Might the same phenomenon occur in other situations and, if so, under what conditions? For such processes to affect trait evolution they would have to change the parameters like population size, genetic variance, selection intensity, or founder effects. Drift processes dominate when population sizes are small, and those processes produce random Brownian-motion style trait evolution; selection, including selective sweeps that affect many genes, dominates when population sizes are large (e.g., Wright 1931; Lande 1976; Wood et al. 2016). Compressing or expanding the geographic range of a species, fragmenting it or allowing fragments to coalesce, or applying widespread selection to cope with an altered environment would be the logical mechanisms for an Earth-system process to impact trait evolution. To affect the sedimentary record that same process would have to alter weathering, transport, accommodation space, or erosion. Climatic, tectonic, and sea-level processes have obvious links to weathering, transport, rates of burial, and accommodation, although so too might any process that controls vegetation cover, karstification, or subsidence (e.g., Valentine 1973; Chakrapani 2005; Katz 2005; Catuneanu 2006; Jeffrey et al. 2016).

Genetic surfing was recognized from molecular phylogeographic analysis of post-glacial expansion of species geographic ranges in North America and Eurasia (e.g., Hewitt 1996; Excoffier and Ray 2008) and later shown to affect trait evolution as well (e.g., Ledevin et al. 2010; Polly 2019b). The terrestrial vertebrate species in those studies have geographic ranges that are far larger than the patchy sediment traps in which their fossil remains are found, yet there is no question that the same glacial-interglacial climatic processes that cyclically altered their geographic ranges also drove cyclic changes in karstification, weathering and transport, water table levels, and therefore the cave and river terrace deposition that is the source of the paleontological record for most small Quaternary vertebrates (e.g., Schreve and Bridgland 2002; Bartolmomé et al. 2021). Quaternary sea level may have had parallel effects on continental shelf environments (e.g., Valentine and Jablonski 1991). Vrba’s “turnover-pulse” theory of climatic- environmental-evolutionary links in which the driver is interaction between tectonics and Quaternary climate in east Africa (e.g., Vrba 1993) is also an example where Earth-systems may simultaneously affect trait evolution and sedimentary deposition, as are the tectonic uplift examples that drive basin formation, habitat fragmentation, and climatic change (e.g., Badgley et al. 2017; Loughney et al. 2021; Weaver et al. 2024). Ancient changes in sea level driven by tectonics are known to have radically reorganized the biogeography of species, sometimes cyclically, creating additional opportunities for an extended common-cause impact on microevolution of morphological traits (e.g., Stigall 2019).

The common-cause hypothesis suggested that the geosphere and biosphere may have coevolved through macroevolutionary processes. The extended common cause suggests that the links may extend to microevolutionary trait evolutionary processes, especially ones that govern rates of evolution. The multi-rate, multi-mode patterns of evolution that are produced by these Wrightian processes and the potential correlation between rate shifts and fossil preservation potential both suggest that fresh scrutiny be given to the methods we use and the assumptions we bring to the study of evolution in the fossil record.

## Acknowledgements

This paper is dedicated to Jim Valentine, who passed while it was being written. When I was a student, he expanded my thinking through the classes I took from him and taught with him, and he continues to teach me through his books and papers. Andrew Bush, David Fox, Steve Holland, Gene Hunt, Gary Motz, and Phil Novack-Gottshall shared thoughts that helped me formulate my thinking. Jessica Utrup, Eric Lazo-Wasem, Susan Butts, Jessica Cundiff, Lourdes Rojas, Lily Berniker, and Mark Siddall did the same, and helped with specimens. Morgan Hill (American Museum of Natural History) and Brian Bagley (University of Minnesota) oversaw CT scanning of snails, and Matt Allen (IU) and Murat Maga (Seattle Children’s Hospital) provided computational advice. The Yale Institute for Biospheric Studies, an Earth-Life Transitions grant from the US National Science Foundation (EAR-1338298), and the Robert R. Shrock Professorship at Indiana University funded my work. The Indiana University Pervasive Technology Institute and the Lilly Endowment, Inc. provided supercomputing resources to run the computational models. Finally, thank you to Linda Ivany and Don Prothero for inviting me to participate in this volume (as well as for editorial contributions), and to Steve Holland and two anonymous reviewers for suggestions that considerably improved the paper.

## Data Availability Statement

All supplementary files are available at Zenodo at https://doi.org/10.5281/zenodo.11636279.

Model output and geometric morphometric data are available at Dryad at https://doi.org/10.5061/dryad.jh9w0vtf4.

## Code Availability Statement

All supplementary code used in this paper is available at Zenodo at https://doi.org/10.5281/zenodo.11636279

## Supplemental files

Supplemental files and code are available at Zenodo (https://doi.org/10.5281/zenodo.11636279) and supplemental model output and semilandmark data are available at Dryad (https://doi.org/10.5061/dryad.jh9w0vtf4) a

**Supplement 1.** Extended methods, data, and figures.

**Supplement 2.** Mathematica code

**Supplement 3.** Zip file of model outputs, including animated maps of trait evolution.

**Supplement 4.** TPS file of *Poecilozonites* aperture semilandmarks.

